# Infantile amnesia results from differential memory retrieval in humans

**DOI:** 10.64898/2026.10.19.755340

**Authors:** Sarah D. Power, Annika Werwach, Aaron Peikert, Marius Fey, Thalia Forbes-Mitchell, Josefine Hild, Claudia Buss, Ulman Lindenberger, Tomás J. Ryan, Markus Werkle-Bergner

## Abstract

Infantile amnesia, the inability to recall memories from our earliest years, is the most common form of forgetting in mammals^1^. Whether early memories never form, decay, or persist but become inaccessible remains unknown. Here, we investigated the nature of forgetting during the infantile amnesia period by longitudinally testing toddlers aged 18-24 months across the complete memory cycle from encoding, initial retrieval one day later, and delayed retrieval up to six months later, in a novel naturalistic episodic memory paradigm. Computational modelling revealed that forgetting during the infantile amnesia period was heterogeneous. Toddlers were as likely to retain the full memory as to lose it completely, while nearly a quarter retained only part of the memory. Encoding accuracy predicted which toddlers retained the full memory at the test one day later, but not at the delayed test up to six months on. Instead, a retrieval-specific EEG signature predicted complete, but not partial, memory retrieval, distinguishing which toddlers had retained the full memory before their behavioural response. Finally, leveraging the COVID-19 pandemic as a natural experiment, we found that maternal immune activation during pregnancy, as in rodents^2^, alters vulnerability to infantile forgetting in humans. Together, these findings show that early memories formed during the infantile amnesia period are not simply erased but persist in an increasingly inaccessible form, a transformation whose trajectory can be shaped by both the neural retrieval mechanism and the prenatal environment.

## Main Text

Humans retain almost no episodic memories from the first years of life ^3,4^. This phenomenon, termed infantile amnesia, is seemingly the most prevalent form of forgetting in the human population ^1^. Three explanations have been posited: early memories may never form (encoding failure), may form but decay (storage failure), or may form and persist in a state that later retrieval cannot access (retrieval failure). Prior evidence shows that infants who later succeed at recalling a learned action sequence show neural evidence of encoding indistinguishable from those who later fail ^5^ and hippocampal activity during encoding predicts subsequent item memory as early as one year of age ^6^, while memory for object-word pairs has been reported at 14-17 months, depending on sleep^7^ . Together these suggest that basic encoding processes are functional in infancy. Yet, whether these early memories are later lost or simply inaccessible cannot be settled by encoding measures alone and has not been tested directly in human development. Rodent studies resolve this, artificially reactivating memory engram cells, labelled during infant learning, reactivates memories that appear permanently lost in adult mice, showing the engram survives even when it cannot be voluntarily retrieved ^2,8^. This capacity is itself developmentally malleable - maternal immune activation during pregnancy prevents the onset of infantile amnesia in male rodent offspring, linking the persistence of early memories to prenatal and postnatal factors ^2,9^. More broadly, prenatal inflammation is a recognised mechanism of developmental programming, maternal inflammation during pregnancy is associated with altered offspring brain development and subsequent cognitive and behavioural outcomes in humans ^10,11^. This raises the possibility that maternal immune activation influences memory persistence not only in rodents but across species.

Despite the mechanistic advances from rodent systems neuroscience, comparable evidence in humans remains lacking. Studies tracking infantile amnesia across a full memory cycle are rare ^12^ and very few systematically trace forgetting curves as a function of age (for exceptions see ^13,14^). Neural correlates that predict which toddlers show forgetting remain unknown. To our knowledge, no study has coupled delay manipulations with neural measures spanning the full memory cycle from encoding to retrieval, a test that is relevant to whether episodic memories in infancy are early expressions of a developing episodic system rather than a separate, transient phenomenon ^15^. Furthermore, no interventional experiments or naturally occurring variation have been used to manipulate infantile amnesia in humans. To bridge these gaps, we built an immersive, naturalistic, multidimensional paradigm to characterise differential memory retrieval across the infantile amnesia period. The task itself requires binding an item, its location, and its surrounding context into a single naturalistic episode where 18-to-24-month-old toddlers are tracked across the complete memory cycle from encoding, through initial retrieval one day later, to delayed retrieval up to six months afterwards. To capture this, we combined top-down video recording of free exploration with analytical tools originally developed for rodent tracking, DeepLabCut ^16^, for markerless pose estimation and Rtrack ^17^ for search-strategy classification, adapted here to toddlers. We recorded mobile EEG in a subset of participants to obtain a neural correlate of memory retrieval. Finally, we leveraged the COVID-19 pandemic as a naturally occurring test of the developmental-programming hypothesis in humans, asking whether prenatal maternal immune activation shapes memory persistence in humans as it does in rodents ^2^.

### Encoding is robust from 18 months

To study forgetting across the complete memory cycle from encoding to retrieval, we tested memory after extended delays in a naturalistic paradigm, binding item, location, and context into a single episode ^12,18^, tapping into real-life events as closely as possible. First, 388 toddlers aged 18 to 24 months were tested in a naturalistic association memory paradigm, in which they encoded the location of a toy character, which was the same toy across contexts but hidden in a different location within each of two distinct visual contextual environments (an underwater and a jungle scene) (Fig. 1A). All stimuli, the toy character and visual contexts, were novel and existed only within the laboratory, ensuring that any subsequent memory for the associations could only derive from the encoding session. Memory retrieval was assessed 1 day after encoding (initial retrieval; IR) and again after a delay of 1, 3, or 6 months (delayed retrieval; DR); exact age and elapsed delay were known to the day for each child. On every trial, the toddler’s first response (the first box they opened) was scored, and each encoding trial continued until they found the toy. Responses were classified as correct context-conditional item-location binding (accurate), systematic context-incorrect (contextual error), or random (random error) (Fig. 1B). On average, toddlers learnt the associations over the course of encoding, with accurate first responses increasing and errors declining across the three trials within each context (Context A and Context B, both P < 0.001) at equivalent rates for the two contexts (trial × context interaction, P = 0.316) (Fig. 1B). The context encountered first also did not influence learning rate or overall accuracy (trial × first context interaction, P = 0.898; main effect of first context, P = 0.258). Older toddlers completed more encoding trials (b = 0.62, t(386) = 6.21, P < 0.001) (Fig. 1C), yet additional trials did not produce more accurate encoding (b = -0.004, t(385) = -1.18, P = 0.239). Encoding accuracy was above chance (P < 0.001) and encoding accuracy did not vary with age at encoding when controlling for the number of encoding trials (b = 0.005, t(385) = 0.67, P = 0.505, R^2^ = 0.004) (Fig. 1D). Learning of the associations was therefore robust and uniform across contexts, and encoding accuracy was independent of age within this range.

**Fig. 1.**
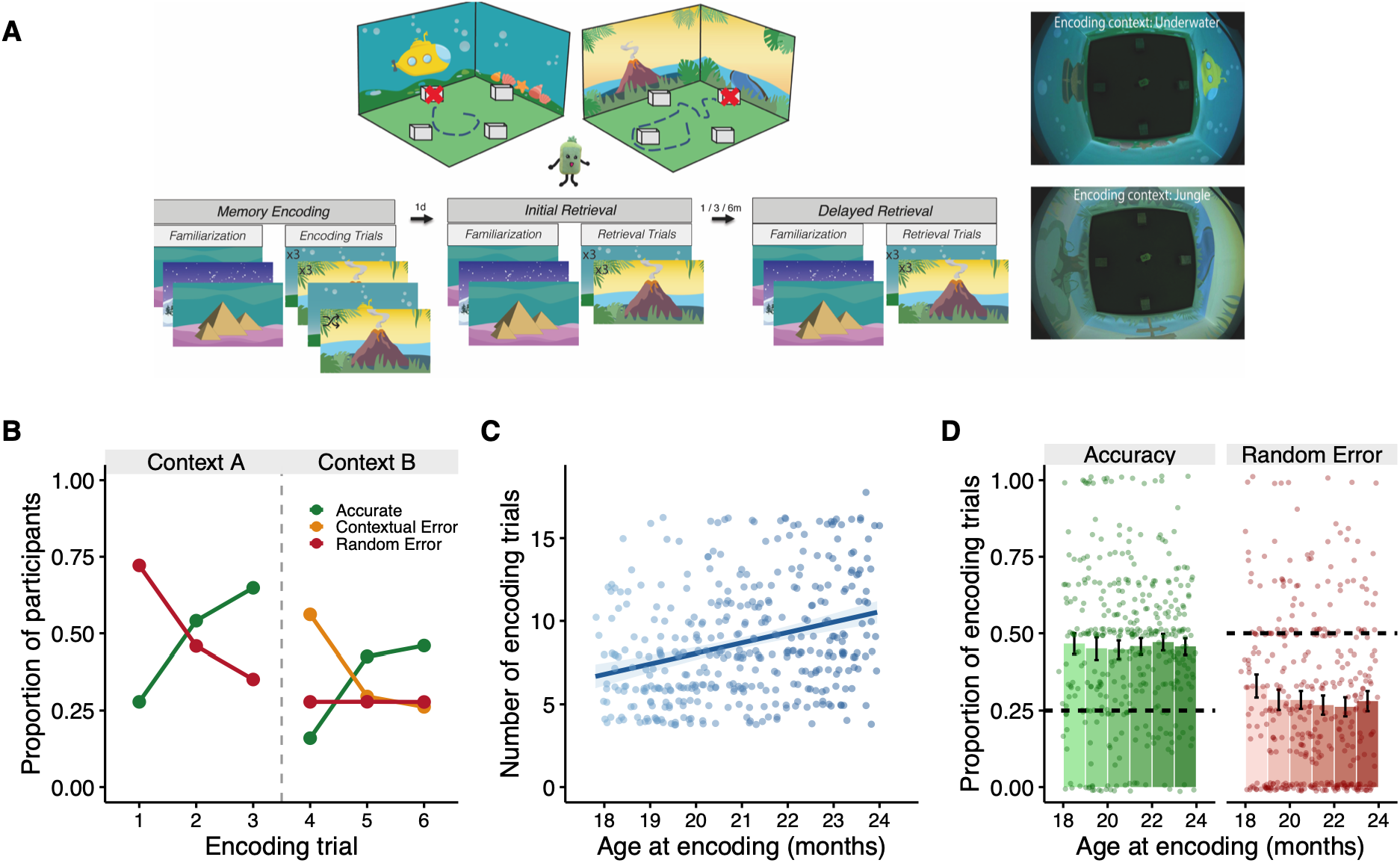
Encoding of item-context-location associations is robust from 18 months. (**A**) Schematic of the behavioural paradigm. (**B**) Proportion of toddlers whose first response on each trial was accurate (green), a contextual error (orange) or a random error (red), across the first 6 encoding trials. Trials 1-3 were presented in context A and trials 4-6 in context B (dashed line marks the context switch). Contextual errors are only defined from trial 4. (**C**) Number of encoding trials by age at encoding (months). Line, linear fit; shading, 95% confidence interval (**D**) Encoding performance by age at encoding (months), shown separately for accuracy (proportion of accurate first choices) and random errors. Bars show the mean and error bars the standard error of the mean (SEM), points show individual toddlers (jittered), dashed horizontal lines indicate chance (0.25 for accuracy and 0.50 for random errors). In (C) and (D) each point is one toddler. N = 388 (18 months, n = 56; 19 months, n = 69; 20 months, n = 71; 21 months, n = 62; 22 months, n = 71; 23 months, n = 59.

### Toddlers rapidly learn to navigate

To characterise the behavioural process underlying encoding, we applied markerless pose estimation (DeepLabCut ^16^) and automated spatial search strategy classification (Rtrack ^17^) (Fig. 2A), tools originally developed for rodent behavioural neuroscience, to toddlers’ movements during the encoding session. Toddlers’ search strategy shifted toward goal-directed navigation over the course of encoding where the probability of an allocentric strategy rose sharply across trials within Context A (β = 0.90, z = 7.90, P < 0.001) (Fig. 2B); individual strategy types in Extended Data Fig. 1A). This dropped sharply at the start of Context B relative to the start of Context A (β = 1.12, z = 4.71, P < 0.001). This drop was accompanied by a sharp rise in contextual errors (perseverance toward the previous, Context A goal location), which became the dominant response at the first Context B trial. A higher proportion of toddlers showed contextual errors (50.7%) than allocentric (21.2%) or non-goal-oriented (28.1%) responses, indicating that toddlers continued searching at the location that had been correct in Context A, rather than reverting to unguided search. However, the rate of improvement across trials within Context B differed from that within Context A (trial × context interaction, P = 0.015). Allocentric use recovered quickly after the first trial but levelled off below Context A levels, indicating that the underlying skill, though momentarily misapplied, largely transferred across the context switch. Once context presentation was randomised across trials (trial 7 onward), further gains largely plateaued (slope reduction relative to Context A (β = -0.89, z = -7.50, P < 0.001), consistent with strategy use approaching ceiling. Movement efficiency followed the same structure (Fig. 2C, D). Both path length (β = - 0.67 m/trial, P < 0.001) and total time (β = -29 s/trial, P < 0.001) decreased steeply across Context A trials, then showed progressively shallower gains across Context B and the randomised block (all P < 0.001), mirroring the strategy pattern above. The movement-derived strategies were behaviourally meaningful. Allocentric strategy use was strongly associated with same-session encoding accuracy (β = 0.46, r = 0.45, P < 0.001) (Extended Data Fig. 1B). This close coupling with encoding accuracy suggests that search strategy indexes, at least in part, memory encoding of item-context-location associations in toddlers 18-24 months.

**Fig. 2.**
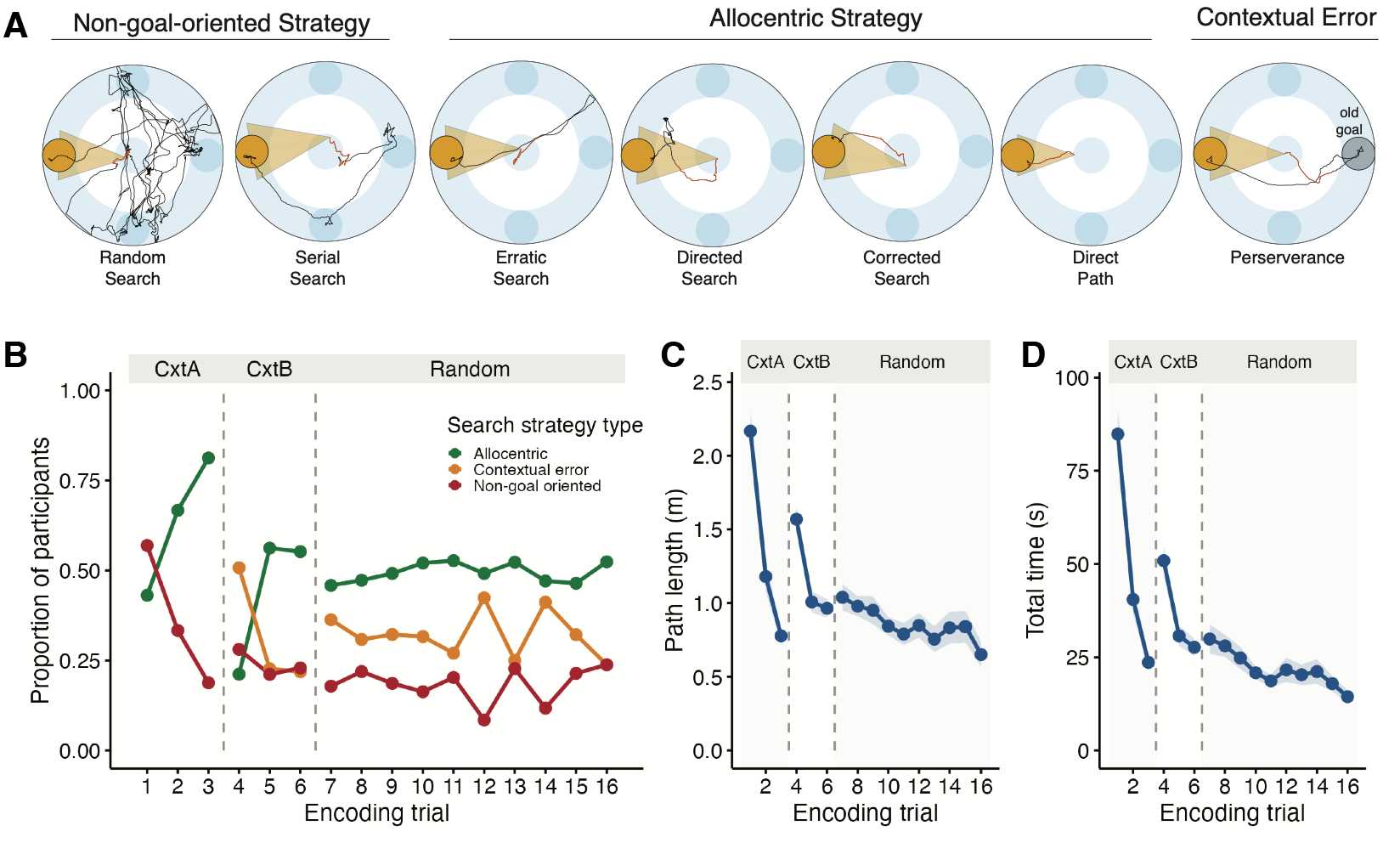
Toddlers shift toward allocentric search strategies over the course of encoding. (**A**) Examples of algorithm-based search strategy classification (Rtrack) in toddlers from this study. Classified strategies are collapsed into three categories: allocentric (direct path, corrected path, directed search, erratic search), non-goal-oriented (random, serial), and contextual error (perseverance toward the previous goal location). (**B**) Proportion of toddlers classified into each strategy category across encoding trials. (**C**) Path length and (**D**) total time per encoding trial, lines show the mean and shading the SEM. Trials 1-3 were presented in context A (CxtA), trials 4-6 in context B (CxtB), and trials 7-16 with randomised context presentation (random); dashed lines mark the context transitions. N = 270.

### Memories survive months-long delays

We next asked whether toddlers can retain these memories and what determines memory retention. One day after encoding, toddlers on average retrieved the associations above chance (P < 0.001) and as at encoding, IR accuracy did not vary with age at encoding (Fig. 3A). Encoding performance tracked IR performance for both accuracy (r = 0.22) and random errors (r = 0.35; both P < 0.001) (Fig. 3B). Across the delay, overall average memory retention was largely stable in every age group (Fig. 3C), even the 18-20 month old group showed robust memory retention up to 6 months later (P < 0.001). However, individual differences were present, with some toddlers showing memory retention, while others forgot substantially (faint lines in Fig. 3C). Linear mixed models do not reveal evidence of strong forgetting on the group level (session, P = 0.330; session × age, P = 0.576), despite big interindividual differences in change. We also regressed the change in accuracy (IR - DR) on candidate predictors while adjusting for IR accuracy. Retrieval delay, age, sex, number of encoding trials, and encoding accuracy were each unrelated to memory retention in this sample (smallest P = 0.110) (Fig. 3D). Group-level accuracy therefore showed stable retention alongside marked individual differences that averages of this kind cannot explain.

**Fig. 3.**
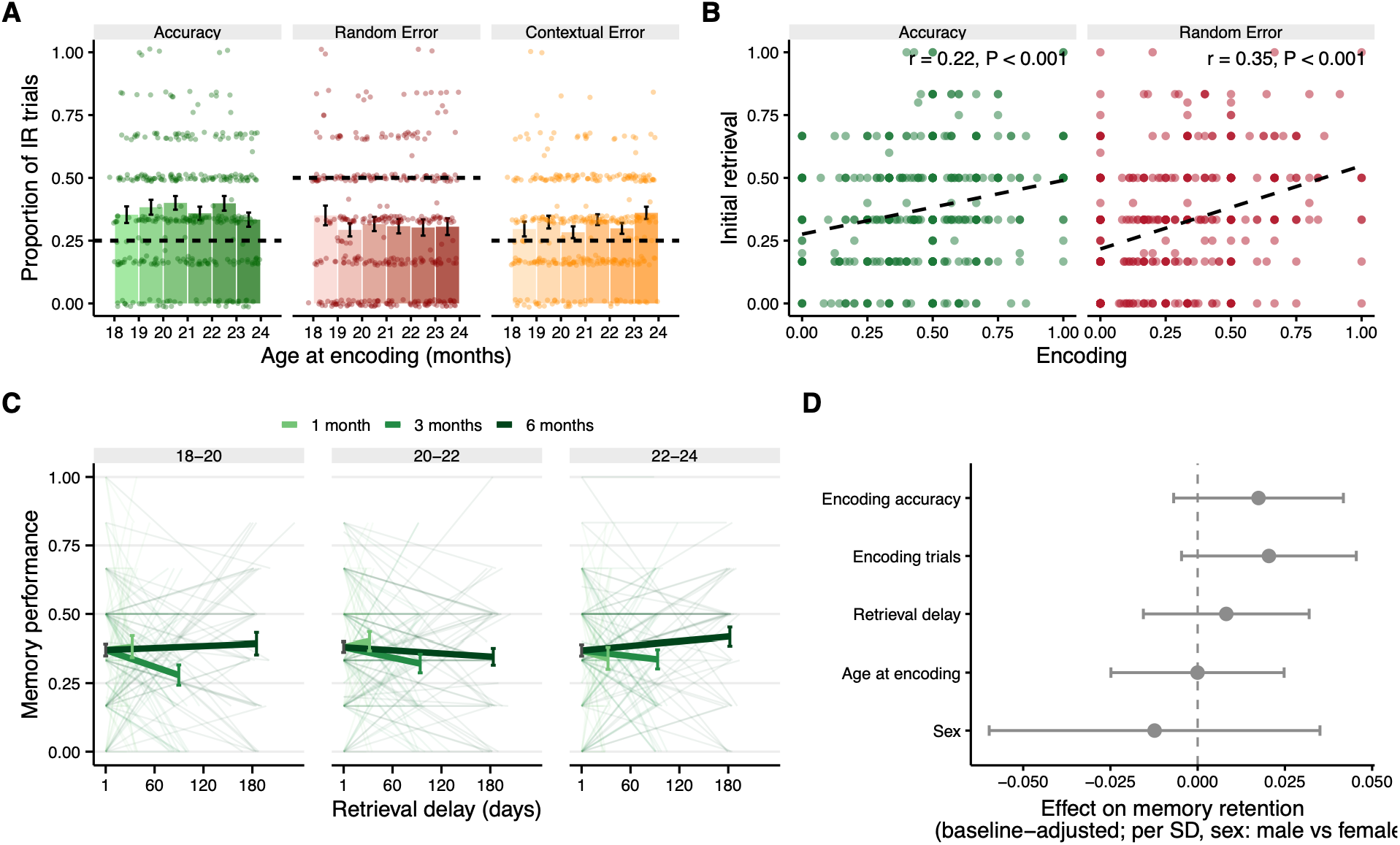
Toddlers as young as 18 months demonstrate long-term memory retention. (**A**) IR performance (1 day after encoding), shown separately for correct responses (accuracy), random errors, and contextual errors. Bars show the mean per one-month age bin, and error bars the SEM; points show individual toddlers plotted at their exact age, dashed lines indicate chance. (**B**) Association between encoding and IR performance; points are individual toddlers and dashed lines the linear fit. (**C**) Associative-memory accuracy across the exact retrieval delay (days), from IR to DR, faceted by age at encoding and coloured by delay condition (1, 3, or 6 months; light to dark; n=122, 130 and 136 toddlers). Faint lines show individual toddlers; bold lines show group means, with error bars (SEM) at the shared initial mean and at each group’s delayed mean. (**D**) Standardised regression coefficients (points) with 95% confidence intervals (bars) from a baseline-adjusted linear model predicting memory retention (IR-DR) from encoding accuracy, number of encoding trials, retrieval delay, age at encoding, and sex, adjusting for IR accuracy; the dashed line marks no effect; two-sided t-tests. N = 388.

Associative memory is often assessed indirectly through recognition of a single item or its context ^12^; we tested whether these simpler measures could account for performance on the associative task itself. In our task, toddlers were tested for item (toy character) and context recognition before entering the testing space, both at IR and DR. Although most toddlers correctly identified the toy at IR (88.7%), toy recall declined with delay (χ^2^(1) = 34.31, P < 0.001) yet did not independently predict associative retention (β = 0.001, P = 0.983), indicating that item and item-context-location memory make independent contributions to retention (Extended Data Fig. 2). Context recall differed markedly by scene at IR, toddlers recalled the underwater scene far more often than the jungle scene (88% vs. 35%; McNemar χ^2^(1) = 131.22, P < 0.001), yet whether toddlers remembered neither, one, or both scenes had no bearing on associative accuracy at IR or DR, indicating that memory for item-context-location associations were independent of memory for the context alone (Extended Data Fig. 3). Vocabulary (age- and sex-normalised, assessed at encoding) did not predict encoding accuracy, IR accuracy, DR accuracy, or memory retention (all P > 0.71), indicating that language ability was not a confound of these results (Extended Data Fig. 4). Neither encoding fidelity, elapsed time, age, single-feature item or context memory or vocabulary emerged as a determinant of memory retention.

### Hidden Markov modelling of memory performance reveals the nature of associative forgetting

These accuracy-based analyses describe forgetting as a single continuous quantity and cannot distinguish partial from complete memory loss. Since episodic-like memory in this task is defined by the joint binding of item, location, and context, rather than by any one of these elements alone, a child can retain parts of this structure while forgetting others. Plain accuracy, which scores only whether the correct location was found, cannot distinguish a child who retains the full item-context-location binding from one who retains only a location stripped of its context. To account for their joint structure, we fit a Bayesian Hidden Markov model (HMM) with seven latent memory classes to each toddler’s trial-by-trial responses, exploiting the sequential structure of the task, trial order and the boundaries between encoding, IR, and DR, to identify discrete patterns of performance based on the joint distribution of accurate responses, contextual errors, and random errors. The model distinguishes seven response classes forming a graded hierarchy of context-conditional binding (Table 1).

**Table 1.**
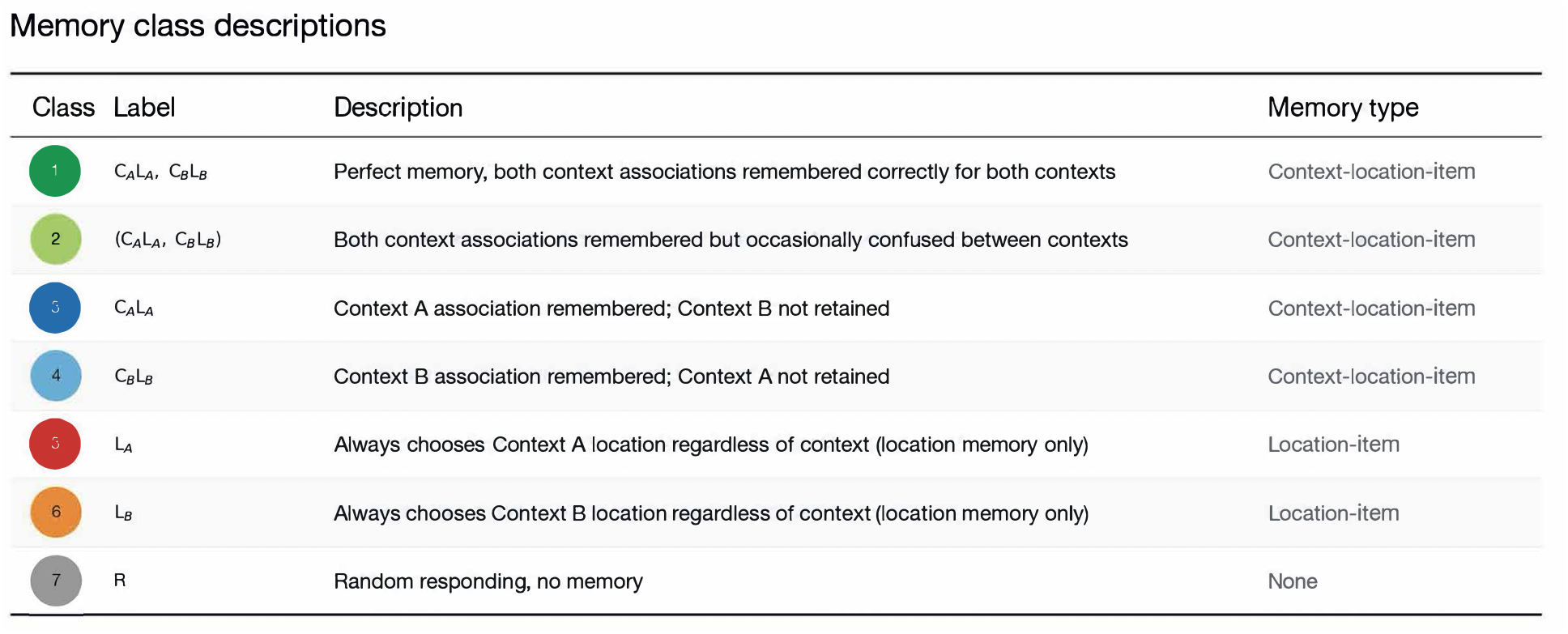
Hidden Markov model memory class descriptions. The seven memory response classes postulated by theory, forming a graded hierarchy of context-conditional binding. Two classes reflect intact binding of both context-location associations (Perfect, class 1; Both associations but imperfect, class 2); two reflect single-context binding, in which one context-location association is retained and the other lost (Context A, class 3; Context B, class 4); two reflect location-only responding, in which a child reliably chooses one location irrespective of context (Location A, class 5; Location B, class 6); and one reflects random responding with no retained memory (class 7). The HMM analysis was used to probabilistically assign individuals to classes based on their actual behaviour.

| Class | Label | Description | Memory type |
| --- | --- | --- | --- |
| 1 | $C_A L_A, C_B L_B$ | Perfect memory, both context associations remembered correctly for both contexts | Context-location-item |
| 2 | $(C_A L_A, C_B L_B)$ | Both context associations remembered but occasionally confused between contexts | Context-location-item |
| 3 | $C_A L_A$ | Context A association remembered; Context B not retained | Context-location-item |
| 4 | $C_B L_B$ | Context B association remembered; Context A not retained | Context-location-item |
| 5 | $L_A$ | Always chooses Context A location regardless of context (location memory only) | Location-item |
| 6 | $L_B$ | Always chooses Context B location regardless of context (location memory only) | Location-item |
| 7 | R | Random responding, no memory | None |

At the end of encoding, toddlers’ memory classes clustered at two poles; around half had bound both context-location associations, and a smaller proportion were responding at random, with location-only responding comparatively rare (Fig. 4B). That sharp separation, however, did not survive to the first retrieval test. By IR, the two-class pattern had already given way to a full spectrum of partial memory. Intact binding remained present in 31% of the sample (split across perfect, both-associations, and single-context forms), location-only responding accounted for 26%, and 43% showed no reliable memory. The subsequent retrieval delay pushed this distribution further toward loss, though considerably more modestly than the 1-day retrieval test had. Intact binding fell only slightly further by DR (29%), random responding grew to become the largest single class (50%), while location-only responding held comparatively steady (26% to 20%). To investigate the nature of forgetting over the delay, we followed toddlers who had bound both associations at IR through to the delayed test. Weighting each child by their probability of having bound both associations at IR, we found that forgetting was not all-or-nothing. Toddlers either lost the full binding completely (45.0%), retained it (32.3%) or slid partway to location-only memory (22.7%). Toddlers in the last group kept where the toy was hidden but not which context it belonged to, a partial state of memory that a single accuracy score does not capture.

**Fig. 4.**
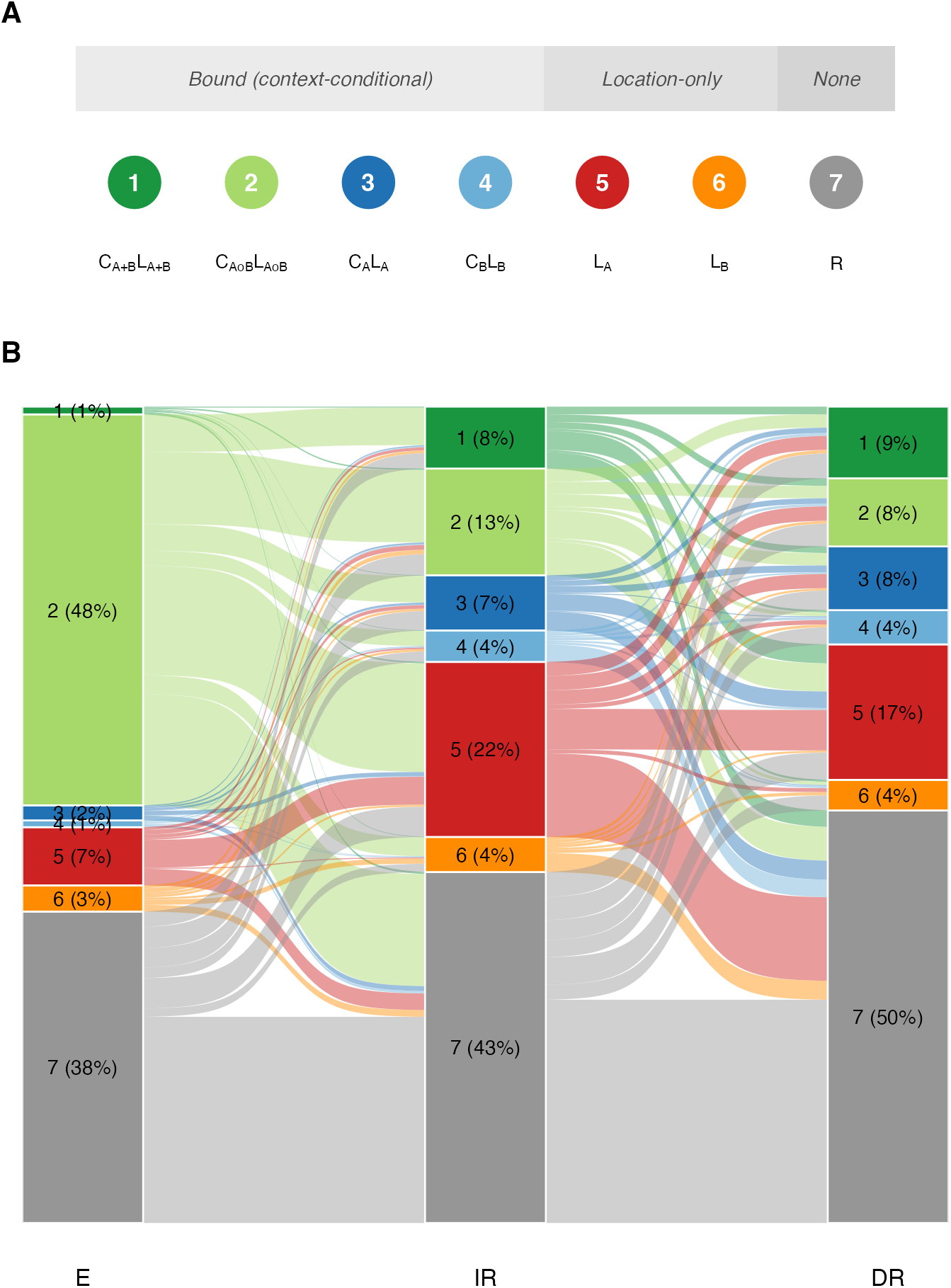
Latent memory classes show graded, not all-or-none, associative forgetting. (**A**) The seven memory response classes forming a graded hierarchy of context-conditional binding. (**B**) Alluvial diagram tracking each toddler’s class membership assignment across encoding (E), initial retrieval (IR), and delayed retrieval (DR). Both flow widths and stratum heights are probability-weighted. Each toddler contributes to every class in proportion to their posterior probability of membership, rather than being assigned to a single most-likely class. Percentages give the probability-weighted share of toddlers in each class at each phase. N = 388.

We next asked what determined which toddlers’ associative bindings dissolved into fully random responding across the delay, the clearest-defined endpoint for complete forgetting, versus which showed graded partial retention. We modelled each toddler’s continuous posterior probability of complete forgetting (class 7), full binding, and location-only responding as a function of age, encoding trials, encoding accuracy, and retrieval delay, with retrieval timepoint (IR vs. DR) and its interaction with each predictor entered in a single mixed model per outcome with a random intercept per child. Encoding accuracy and encoding trials were the strongest predictors of complete forgetting (F(1,383) = 48.19 and 16.97, both P < 0.001). Encoding accuracy’s protective effect was substantially weakened at DR (accuracy × timepoint: F(1,383) = 22.06, P < 0.001; simple slopes on P(Random): IR = -0.52 vs. DR = -0.15), whereas the trials × timepoint interaction was not significant. P(Random) itself rose significantly from IR to DR (F(1,383) = 10.02, P = 0.002), confirming that complete forgetting continues to accrue between IR and DR. A similar pattern held for full binding retention (accuracy P < 0.001 and trials P = 0.004; accuracy × timepoint, P = 0.002; trials × timepoint, P = 0.023, adjusted P = 0.051) and, for encoding accuracy, for location-only responding (timepoint interaction, P = 0.018, adjusted P = 0.043); location-only responding additionally showed continued decay with longer delays that the other two outcomes did not (timepoint × delay, P = 0.023, adjusted P = 0.043). Age predicted none of the three outcomes at either timepoint. Together, these results show that encoding quality strongly predicts how much toddlers remember one day later. Still, its predictive power fades by the delayed test and does not determine which discrete memory class (fully intact or partial) an toddler ends up in.

### Neural correlates of episodic retrieval in toddlers

We next turned to retrieval itself, asking whether a neural measure obtained at test could distinguish toddlers who went on to succeed from those who did not. In a sub-cohort of the sample, 32-channel mobile EEG was recorded during the memory task in toddlers aged 18-20 months with a DR interval of 1 or 3 months. To identify neural correlates of memory performance, we applied behavioural partial least squares (PLS) analysis to EEG spectral power (1-25 Hz) recorded during encoding, IR and DR in the 76 participants who contributed EEG data across all three phases. The first latent variable (LV1) was significant (P = 0.0002) and accounted for 82.1% of cross-block covariance (Fig. 5A). Critically, the behavioural salience for LV1 was concentrated in the DR phase (Fig. 5B), indicating that the neural signature is most strongly expressed during retrieval after an extended delay. The brain-behaviour correlation was positive (r = 0.17, P = 0.009), indicating that participants with higher retrieval accuracy expressed the associated neural pattern more strongly, predicting rather than following correct behavioural responding (Fig. 5C). Inspection of bootstrap ratio scores (BSR > 2.5) revealed two spatially and spectrally distinct reliable contributions to LV1 (Fig. 5D). Low-frequency power (<5 Hz) over parieto-central regions and higher-frequency power (>12 Hz) over fronto-central regions were both expressed more strongly in participants with higher retrieval accuracy. To determine whether the DR LV1 brain score was associated with the specific pattern of associative memory, we modelled DR LV1 as a function of each toddler’s posterior probability of full context-location binding (summing the perfect, both-associations, and single-context classes) and probability of location-only responding, entered as continuous predictors in a single model. The probability of full binding was positively associated with LV1 (β = 68.7, P = 0.003), while the probability of location-only responding showed no relationship (β = 10.0, P = 0.559), and the two coefficients differed from each other (P = 0.023). This specificity, a positive relationship with full associative binding, and no relationship with location memory alone, converges with the latent memory analysis in Fig. 4 in distinguishing toddlers who retained the complete episodic trace from those who retained only partial information.

**Fig. 5.**
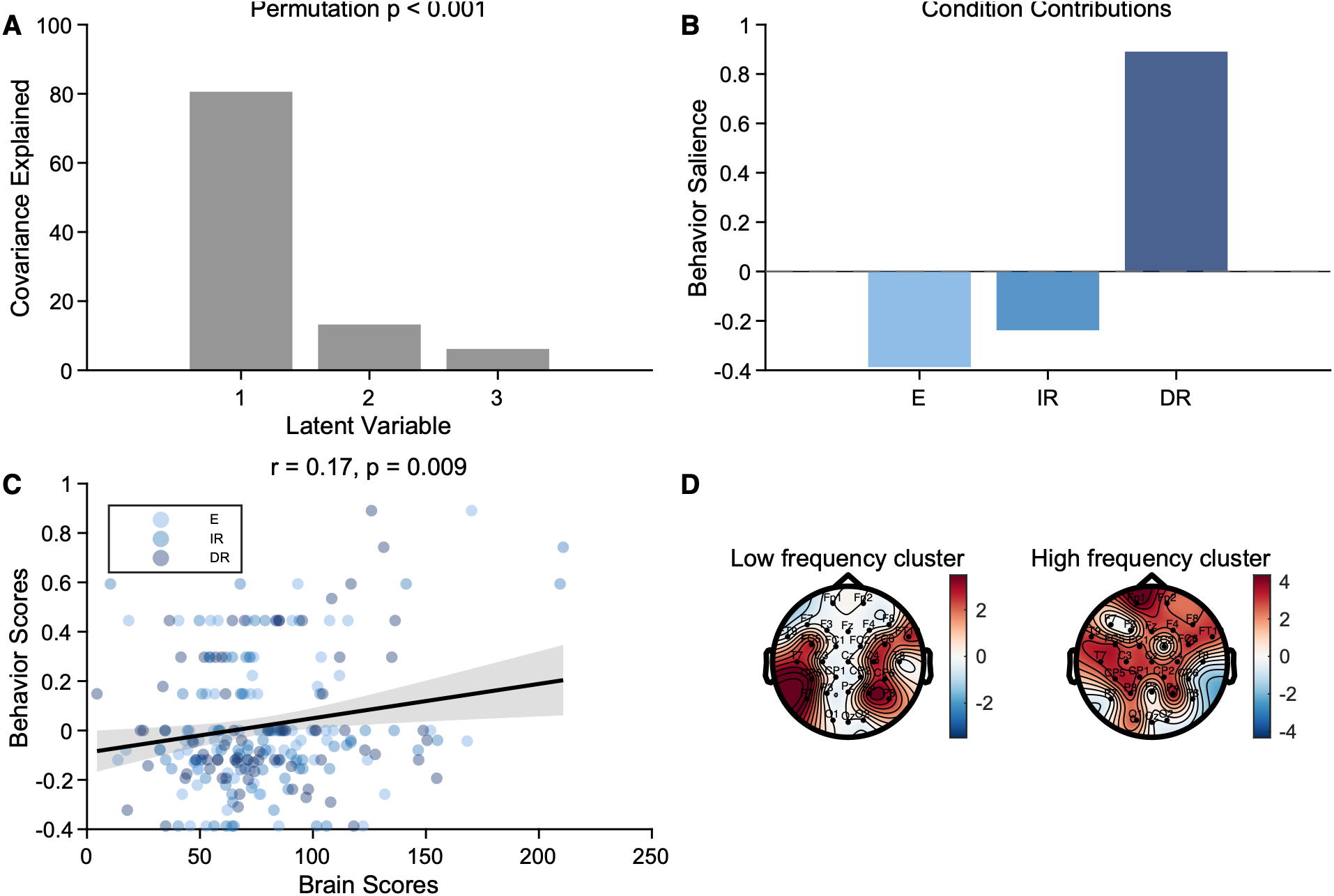
An EEG signature of retrieval distinguishes toddlers who retained the full memory. (**A**) Cross-block covariance explained by each latent variable (LV) from the behavioural partial least squares (PLS) analysis of EEG spectral power (1-25 Hz) across encoding (E), initial retrieval (IR) and delayed retrieval (DR). LV1 explained 82.1% of the cross-block covariance (permutation test, P = 0.0002). (**B**) Behavioural salience of LV1 by session (E, IR, DR). (**C**) Correlation between each toddler’s LV1 brain score and behavioural accuracy score (r = 0.17, P = 0.009). (**D**) Topographical distribution of bootstrap ratios (BSR) for LV1, thresholded at |BSR| > 2.5: a low-frequency (<5 Hz) cluster over parieto-central electrodes and a higher-frequency (>12 Hz) cluster over fronto-central electrodes. N = 76 toddlers (EEG sub-cohort) contributing data at all three phases, each point in (C) is one toddler at one phase (E, IR or DR).

### Prenatal immune activation alters toddler memory performance

No study thus far has used an intervention or naturally occurring variation to manipulate infantile amnesia in humans. We took advantage of a naturally occurring experiment, the COVID-19 pandemic, which exposed a subset of toddlers prenatally to potential prenatal immune activation without any experimental intervention. This allowed us to ask whether the prenatal environment shapes vulnerability to infant forgetting. Toddlers (18-24m) exposed in utero to maternal COVID infection (MCI) (COVID+; n = 90) and age-matched controls from our sample (COVID-; n = 136) completed the associative memory task at encoding, IR and 6-month DR. Both groups learned the associations across fixed encoding blocks (Context A, trials 1-3; Context B, trials 4-6), with equivalent overall encoding performance (Fig. 6A, B). At retrieval, MCI-exposed toddlers showed lower associative accuracy and higher random error across IR and DR (accuracy: b = -0.073, adjusted P = 0.003; random error: b = +0.068, adjusted P = 0.011) (Fig. 6C-E). The number of encoding trials completed predicted 6-month memory retention differentially by cohort (trials × cohort: b = -0.029, P = 0.002) (Fig. 6F). MCI-exposed toddlers who completed more trials showed worse memory retention (b = -0.018, adjusted P = 0.039), whereas the slope in non-MCI-exposed toddlers was not significant (b = +0.011, adjusted P = 0.053). Because trials beyond the sixth require randomised context presentation, these data suggest that MCI-exposed toddlers who completed encoding under increased cognitive load showed accelerated forgetting over a 6-month delay.

**Fig. 6.**
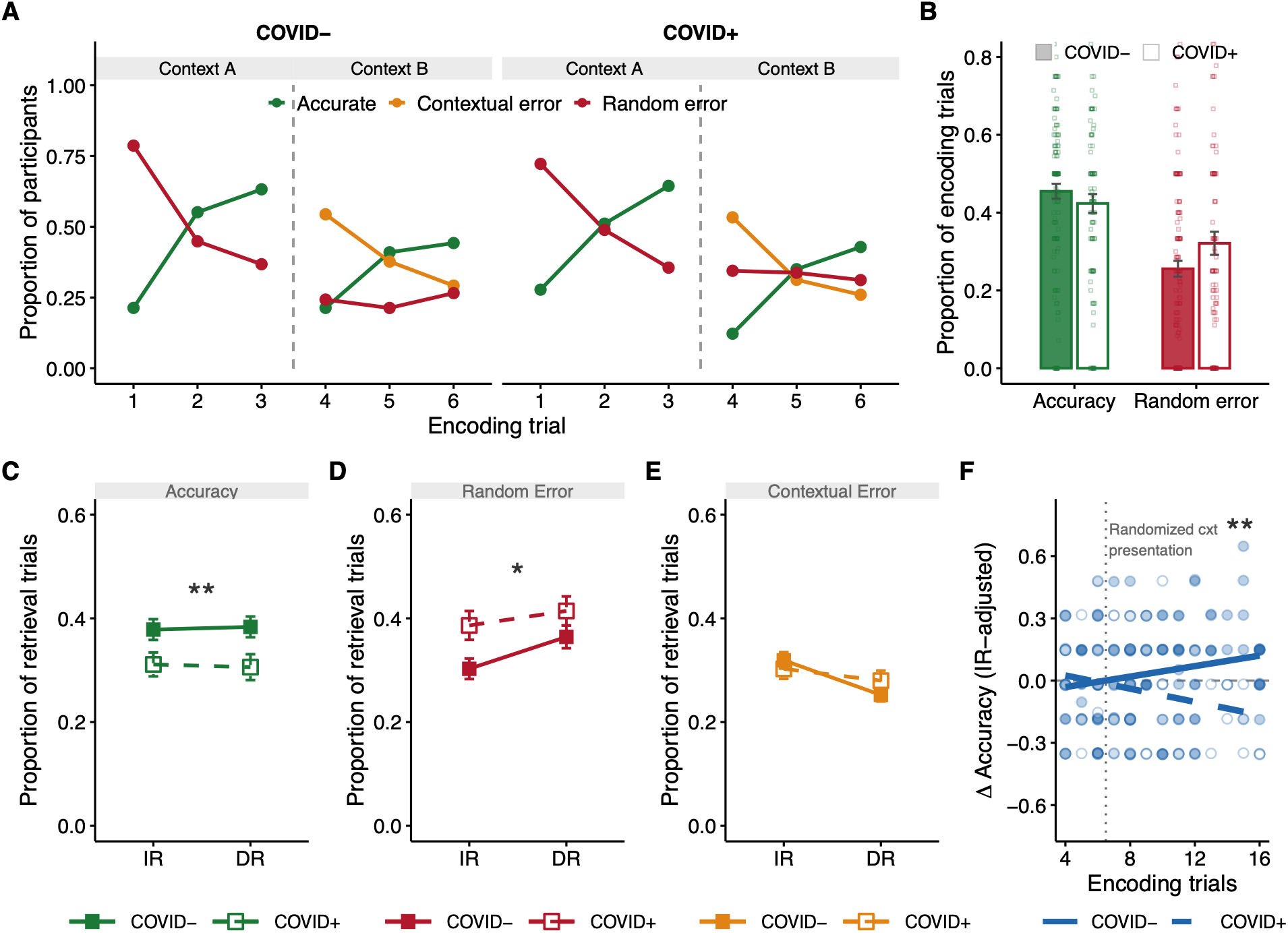
Maternal COVID-19 infection during pregnancy is associated with altered memory in toddlers. (**A**) Proportion of participants giving each response type (accurate, contextual error, random error) across the six encoding trials, separately for COVID- (n = 136) and COVID+ (n = 90) toddlers. Context A and B blocks are marked above. (**B**) Overall accuracy and random-error proportions at encoding, averaged across all encoding trials, by cohort (filled bars, COVID-; open bars, COVID+). Bars show the mean and error bars the SEM, points show individual toddlers (**C-E**) Accuracy, random error, and contextual error at IR and DR by cohort (mean and SEM). Asterisks denote Benjamini-Hochberg-adjusted P values for the cohort effect from linear mixed-effects models (likelihood-ratio test): *P < 0.05, **P < 0.01. (**F**) Baseline-adjusted change in accuracy (IR to DR residual after regressing on IR accuracy) as a function of the number of encoding trials completed by cohort, lines show linear fits and points individual toddlers. The dotted line marks the sixth trial, beyond which context presentation was randomised. **P < 0.01, trials × cohort interaction. N = 136 (COVID-), 90 (COVID+).

## Discussion

A prevailing assumption has been that infantile amnesia reflects an inability of the immature hippocampus to establish durable episodic memories ^19^. Our findings challenge this view by demonstrating differential retrieval strategies across the timespan of infant forgetting. Before the second year of life, toddlers formed stable, context-bound episodic memories independent of age at encoding, yet these memories undergo systematic transformations over subsequent months that progressively reduce their accessibility. By combining HMM with neural measures of retrieval, we show that early toddler forgetting is better explained by changes in the use of memory representations than by overall failures of encoding or storage. This framework allows us to begin to reconcile long-standing inconsistencies in the infant memory literature and suggests that infantile amnesia reflects a dynamic process of representational reorganisation rather than wholesale erasure of early experience.

We found that encoding of complex item-location-context associations was robust and age-invariant. Performance at 18 months, the youngest age tested, was indistinguishable from that of 23-month-olds, with no age gradient in encoding fidelity, whereas earlier work reported a marked improvement in episodic-like binding at about 20-21 months ^18^. This argues against the dominant account that hippocampal circuitry supporting episodic memory remains functionally immature during infancy, rendering encoding unreliable. The capacity to form context-specific episodic-like bound representations is in place before the offset of the infantile amnesia period, and later forgetting cannot be attributed to a failure to encode. This is consistent with rodent evidence that hippocampal-dependent memories acquired during the infantile amnesia period are encoded and stored, but only later expressed once appropriately reinstated ^20,21^. Nor is it merely item memory that is retained, recall of the toy’s character and of the encoding context varied independently of associative accuracy at every stage, indicating that single-feature recognition is neither necessary nor sufficient to explain performance on the associative task. This suggests that paradigms relying on item recognition alone, while valuable, likely underestimate the sophistication of early relational memory as originally defined ^22,23^. Storage deficit accounts of infantile amnesia predict that elapsed time should erode memory. We found that delay length did not predict whether an established binding survived intact. Rather, encoding fidelity, not time, determined whether a memory was retained shortly after learning, though not which infants kept it months later, suggesting that near-term formation and long-term maintenance are governed by partially distinct processes. Preverbal-to-verbal transition ^24^ is also excluded as an explanation of forgetting. Expressive vocabulary, measured independently, did not predict any memory metric, and robust long-term memory was evident even in toddlers with limited productive vocabulary.

Instead, our findings support the view that early memories persist through the infantile amnesia period but progressively become less accessible to retrieval rather than being lost altogether. The HMM analysis provides insight into how this transformation unfolds. Immediately after encoding, memory was represented in an essentially all-or-none manner. Toddlers could either successfully retrieve both item-context-location associations or showed little evidence of memory, with almost no intermediate classes. This pattern is consistent with hippocampal pattern-completion dynamics underlying relational binding ^25,26^. However, this sharp separation had already begun to transform after 24 h, and over delays of several months the distribution shifted further toward memory loss. Importantly, forgetting split toddlers into three outcomes rather than following one dominant pattern. Toddlers were more likely to lose the full memory completely than to retain it, while nearly a quarter instead transitioned into an intermediate representational state in which some, but not all, associations remained accessible, a pattern invisible to conventional accuracy measures but revealed by the HMM. Two additional observations corroborate this claim. First, encoding accuracy predicted how much information toddlers ultimately retained, but its influence on representational class weakened substantially, rather than persisting unchanged, between initial and delayed retrieval. By delayed retrieval, it no longer predicted whether a memory reached full binding, though it still predicted, more weakly, whether it collapsed into complete forgetting. Initial encoding strength therefore appears to determine the quantity of information preserved, whereas the long-term structural transformation of the memory follows a partially independent process. Second, fully bound episodic memories at delayed retrieval were observed across the entire age range studied. Indeed, more than a third of the toddlers classified into the fully bound memory class after the delay were between 18 and 20 months of age, indicating that episodic-like representations are already present well before the offset of the infantile amnesia period.

These findings are consistent with the framework proposed by Ryan and Frankland ^1^. They argue that forgetting more broadly reflects reduced engram accessibility rather than engram loss and propose that infantile amnesia specifically results from developmental circuit remodelling reducing that accessibility while the underlying memory engram persists. Our HMM analysis suggests one possible computational realisation of this process. Rather than fading uniformly, episodic memories appear to transition into qualitatively different representational states whose remaining content differs substantially across individuals. This interpretation is consistent with memory accessibility declining even as some associative content remains behaviourally expressible. The marked variability in these trajectories also has important methodological implications. If different toddlers retain different components of the same experience, standardised paradigms using relatively simple stimuli or limited retrieval cues, while valuable, may fail to capture the accessible information in many participants. Such tasks may therefore underestimate memory retention and contribute to the inconsistent findings that characterise the developmental memory literature.

The EEG results provide converging evidence that the HMM memory classes reflect genuine differences in memory representations. A single latent variable, accounting for 82.1% of the covariance between EEG and behaviour, captured a neural pattern associated with correct responses before the actual behavioural response occurred, indicating that it reflects the neural processes supporting retrieval rather than its consequences. The pattern combined increased power over parieto-central regions in the toddler theta band (3-5 Hz) ^27^, linked to hippocampal-cortical communication during retrieval ^28–30^, and increased power over fronto-central regions above 12 Hz, linked to post-retrieval processing ^31^. This pattern was most strongly expressed during delayed retrieval. This selective emergence of a fronto-central, cortically linked signature at delayed retrieval is consistent with recent causal evidence in rodents that prefrontal and entorhinal engagement in object-location memory recall increases with memory age and becomes functionally necessary specifically for remote, rather than recent, retrieval ^32^. This pattern is in line with broader theoretical accounts in which retrieval of older memories draws increasingly on cortical circuits beyond the hippocampus ^33^. The signature was selectively associated with retrieving fully bound episodic memories, whether it was one or both contexts, rather than location-only memories. Together, these observations suggest that the neural signal indexes the integrity of consolidated episodic representations rather than transient fluctuations in task performance. More broadly, they provide independent support for the central conclusion that developmental forgetting reflects changes in the accessibility of memory representations rather than failures of encoding.

Maternal COVID infection, our natural test of prenatal immune activation, produced a strikingly different pattern than reported in rodents. Whereas MIA abolishes infantile amnesia in male rodent offspring ^2^, we found the opposite here. Although overall encoding was preserved, toddlers born following MCI showed a relationship between learning and retention that differed from controls, where additional encoding trials predicted poorer subsequent memory retention ^34^. This contrast to rodent research may reflect differences in task structure rather than biology. The rodent MIA study tests a single learned association, whereas our paradigm required binding two competing contexts. A single-context design might therefore have revealed a convergent, protective effect instead, as shown in mouse studies. While these findings require replication, they suggest that vulnerability to infantile amnesia may be shaped before birth rather than emerging solely through postnatal maturation. This complements the focus on developmental age with an additional focus on the prenatal factors that influence the long-term stability of episodic memory representations.

## Conclusions

The present findings challenge the long-standing view that infantile amnesia reflects an inability to encode or store episodic-like memories. Instead, they indicate that the capacity to form context-bound episodic-like memories is already established by 18 months, well before the behavioural offset of the traditional infantile amnesia period. Rather than disappearing over time, these memories undergo systematic transformations in accessibility of their representational structure, progressively reducing their accessibility while often preserving substantial elements of the original experience. This framework explains why retrieval success varies across individuals despite comparable encoding, why standard memory paradigms may underestimate early memory, and why behavioural forgetting need not imply that a memory is gone. More broadly, our findings show that infantile amnesia reflects differential memory retrieval in humans, shifting the focus from the existence of early memories to the mechanisms governing their long-term transformation and retrieval in humans.

## Acknowledgments

The research was conducted within the project Lifespan Rhythms of Memory and Cognition (RHYME; PI: M.W.-B.) at the Center for Lifespan Psychology, Max Planck Institute for Human Development, Berlin. We are grateful for the support of all the participating families; all student assistants for their support in recruitment and data collection; past and present members of the RHYME project for scientific discussions and support; and the Scientific Technology & Computing (STC) team at the Max Planck Institute for Human Development for their research infrastructure and technical assistance. We further acknowledge support by the Max Planck Dahlem Campus of Cognition (MPDCC).

