## supplemental file for "Infantile amnesia results from differential memory retrieval in humans"

### Methods

Participants

Data were collected from 534 toddlers aged 18-24 months who completed all three sessions (encoding, immediate retrieval, and delayed retrieval) of an item-context-location memory task. Toddlers were recruited across three overlapping cohorts; one cohort was exclusively COVID-unexposed and completed the task at delays of 1, 3, or 6 months (n=388). A second sub-cohort additionally underwent mobile EEG at all three sessions and included both exposure groups at 1- or 3-month delays (n=76). A third sub-cohort age-matched cohort was exclusively COVID-exposed and completed the task at a 6-month delay (n=90). Analyses characterising typical memory development (Fig. 1-4) use the COVID-unexposed toddlers to establish developmental patterns independent of prenatal exposure. Fig. 5 neural analyses pool across exposure status, as brain-behaviour relationships did not differ by exposure. Fig. 6 directly compares exposure groups, using 226 participants tested at a 6-month delay (136 COVID-unexposed, 90 COVID-exposed); the groups were recruited and tested in the same period and were balanced on age, retrieval delay, number of encoding trials, sex and first encoding context (Supplementary Table 1). Participants were native German speakers with no history of learning disabilities or neurological, psychiatric, or sleep disorders. Participants who did not complete at least four trials in the initial encoding session were excluded from further data collection. The study received approval from the ethics committee of the Max Planck Institute for Human Development (LIP-2022-03/ECHO) and was conducted in accordance with the Declaration of Helsinki. Written informed consent was obtained from parents or legal guardians prior to participation.

Experimental Setup

A new custom-built testing arena (4 x 4 m) was constructed in a controlled laboratory environment. The arena was enclosed by four projection-capable walls displaying dynamic, cartoon-style virtual environments depicting naturalistic scenes (e.g., jungle, underwater) depending on the trial. The arena's entrance was concealed within the projected walls, so that once inside, participants had no visible door or external landmark, and the space appeared fully enclosed. Ceiling-mounted video cameras recorded participant movement. Four identical boxes were positioned at fixed locations in the arena. All visual stimuli, environmental contexts, and the toy figure were uniquely created for the study and were not derived from external sources. This room-scale, immersive setup allowed toddlers to freely explore and search across a real physical space,

Data acquisition:

*Familiarisation:*

Each experimental session began with a familiarisation phase comprising three trials. Participants were exposed to two non-target contexts (magic forest and desert) and instructed to explore all four boxes, confirming they were empty. The toy character was placed in the centre of the arena. Familiarisation procedures were consistent across sessions.

*Encoding:*

During encoding, participants experienced two target contexts (underwater and jungle). Context-location pairings were randomly assigned for each participant and remained fixed across sessions. Participants completed between 4 and 18 trials; the first eight followed a structured order (three underwater, three jungle, one underwater, one jungle), while any remaining trials were randomised. The order and location assignments were counterbalanced across participants. Before each trial, participants interacted with the projected context for 2-3 minutes. Trials began from a central starting point marked by an X following a verbal countdown, with participants starting facing a different direction on each trial. Participants were never shown or told the toy's location in advance; they located it independently on each trial and learned the context-specific association purely through their own repeated search. Participants could open multiple boxes per trial until the toy was found and the trial ended when the correct box was opened. Encoding was limited to one hour.

*Immediate and Delayed Retrieval:*

Memory retrieval was assessed at two time points: an immediate retrieval (IR) session 24 hours after encoding, and a delayed retrieval (DR) session occurring either 1, 3, or 6 months after encoding. Prior to entering the arena at both sessions, participants completed two recognition tasks: (1) item memory (selecting the target toy from five custom figures) and (2) context memory (identifying the two encoding contexts from five images, comprising one familiarisation context and two lures). Participants then completed a maximum of six search trials (three per context). Each trial was preceded by a 2-3 minute contextual interaction and began from the central starting point. Only one box could be opened per trial. The participants did not receive feedback during the search trial.

Markerless pose estimation and navigational strategy analysis

Toddlers movements during encoding were tracked using DeepLabCut v3, PyTorch engine; (*15*), a multi-animal markerless pose estimation toolkit. A multi-animal model was trained to simultaneously track four individuals (toddler, experimenter, and up to two parents) across 17 body keypoints per individual (head, nose, bilateral shoulders, elbows, hands, navel, hips, knees, ankles, and feet) at approximately 30 fps (resolution 1280 x 1440 px). Two model variants were trained on 95% of manually labelled frames: a bottom-up model (shuffle 1; BU; test RMSE = 6.39 px at pcutoff = 0.5) and a crop-then-detect model (shuffle 2; CTD; test RMSE = 2.90 px at pcutoff = 0.5). To ensure detection of the toddler, detections were filtered by shoulder width and limb proportions calibrated to 18-24 month toddlers. Gaps were interpolated using a Kalman smoother. Head position coordinates were extracted per frame and passed to Rtrack (*16*) for search strategy analysis, which classified each trial into one of nine search strategies using a trained random forest classifier based on the Barnes maze model. For a full description of search strategies, see (*16*). Tracked head positions were overlaid on all videos and manually verified to ensure correct toddler identification prior to analysis. Trials for which head position could not be reliably tracked were excluded from analysis.

Language assessment

As a covariate of cognitive development, participants' language abilities were assessed using the FRAKIS-K, a standardised German adaptation of the MacArthur-Bates CDI (*35*). Assessments were completed by parents prior to encoding and delayed retrieval sessions and yielded percentile scores based on vocabulary and grammar norms for 18-30-month-old toddlers.

*Hidden Markov modelling of memory class*

To characterise each toddler's memory class across the three experimental sessions (encoding, initial retrieval, delayed retrieval), we fitted a Bayesian hidden Markov model (HMM) to the trial-by-trial response sequences; full model specification, including prior distributions, transition-matrix structure, and simulation-based parameter recovery, is provided in the appendix (*33*). Based on theoretical considerations (*20*), the model posits seven discrete latent memory classes representing qualitatively different combinations of context and location knowledge: perfect context-and-location memory (class 1), imperfect context memory (class 2), context A only (class 3), context B only (class 4), location A only (class 5), location B only (class 6), and random/no memory (class 7). All toddlers begin in the random class. Emission probabilities are fully determined, except for a single attention parameter ($\alpha \sim \text{Beta}(3,2)$) that captures behavioural noise by mixing each class's theoretical emission probabilities toward the chance-level response distribution expected under random guessing. Six phase-specific theory-informed transition matrices govern class dynamics within and between phases; within-encoding transitions are constrained to be one-directional (no within-session forgetting), encoding-related boundary transitions carry stability priors (Dirichlet, c_diag = 9), and the retrieval-interval (forgetting) boundary uses a flat, uninformative prior with no preference for retention or forgetting. The model was implemented in the Julia programming language and parameters were estimated via the No-U-Turn Sampler (NUTS) with 4000 samples across 32 chains; simulation-based parameter recovery using a reduced three-class instantiation of the model confirmed that it correctly identifies known transition structure. Diagnostics indicated adequate convergence ($\hat{R} < 1.01$, ESS > 100, E-BFMI > 0.2). Per-child posterior class probabilities were obtained using the forward-backward algorithm, yielding filtered and smoothed posteriors; modal class assignments at initial and delayed retrieval were used for downstream statistical analyses.

*EEG recording*

Mobile EEG was recorded in a sub-cohort of toddlers aged 18-20 months with delayed retrieval intervals of 1 or 3 months. Data were recorded from 32 active Ag/AgCl electrodes mounted in an elastic cap (EasyCap) at standard 10-20 positions, with one electrode placed on the left orbital ridge when tolerated by the child. Signals were referenced online to the right mastoid, with AFz serving as ground. Electrode impedances were largely maintained below 30 kΩ. Data were amplified using a wireless LiveAmp system (Brain Products GmbH, Gilching, Germany) and digitized at 500 Hz via Lab Streaming Layer (*36*). Trial events were marked with manually applied triggers, verified against video recordings post-acquisition.

*EEG preprocessing*

Offline preprocessing was performed in MATLAB R2024b and EEGLAB (*37*) and Fieldtrip toolboxes (*38*) with custom scripts (*34*). Data were first high-pass filtered at 1.25 Hz (zero-phase Hamming-windowed sinc FIR), followed by manual removal of severely artifacted segments and bad channels. Independent component analysis was applied, and components were manually classified as neural, ocular, or muscle/movement artefacts. Data were subsequently filtered with a 0.5 Hz high-pass and a 27.5 Hz low-pass (zero-phase Hamming-windowed sinc FIR), and ocular and muscle/movement components were removed. Artefact subspace reconstruction was applied with a cutoff of 10, after which bad channels were spherically interpolated, and data were re-referenced to the average of both mastoids. Trials were visually inspected for residual artifacts and additionally screened for amplitude excursions exceeding ±200 µV in any channel. Participants for whom all trials were rejected were excluded from further analyses. Full details of the preprocessing pipeline, including parameter validation, are provided in (*34*).

*EEG spectral analysis*

Continuous EEG data from each trial, defined as the interval from trial onset until immediately before the participant opened the first box, were segmented into consecutive, non-overlapping 4-s epochs. Spectral decomposition was performed using a multitaper fast Fourier transform approach, yielding an effective spectral smoothing of ±2 Hz around each centre frequency (*39*); power spectra were computed from 1 to 25 Hz in steps of 0.1 Hz. Resulting EEG power spectra were analysed using behavioural partial least squares analysis (*40*, *41*), which identifies multivariate associations between EEG spectral power and behavioural accuracy across experimental phases. EEG power spectra served as the neural measures, and accuracy scores as the behavioural measures, with experimental phase (encoding, initial retrieval, delayed retrieval) modelled as separate conditions, allowing extraction of up to three latent variables (LVs). Statistical significance of each LV was assessed using permutation testing (10,000 permutations), and reliability of electrode/frequency saliences was evaluated using bootstrap resampling (10,000 bootstrap samples). The association between EEG brain score (LV1) and HMM memory class was modelled with linear regression, with each child's posterior probability of full binding and of location-only responding entered as continuous predictors rather than a collapsed categorical assignment.

*Statistical Analysis*

All behavioural statistical analyses were performed in R 4.5.1 using lme4 1.1.37, lmerTest 3.1.3, nnet 7.3.20, MASS 7.3.65 and emmeans 2.0.1. All tests were two-sided with $\alpha = 0.05$. Age was entered as exact age at encoding (days or months), and baseline-adjusted change in accuracy was the residual of delayed minus initial retrieval accuracy after regression on initial retrieval accuracy. Models were fit as ordinary linear or logistic regression (lm, glm) when each participant contributed a single observation, and as linear or generalised linear mixed-effects models (LME/GLMM; lmer, glmer, bobyqa optimiser) when observations were repeated within participants (e.g., multiple trials, timepoints, or response-type metrics per child), with participant as a random intercept; this repeated-measures structure, not the number of predictors, determined the model class, and most models, of either type, included multiple simultaneous predictors. Where a predictor's effect was compared across immediate and delayed retrieval, retrieval timepoint and its interaction with each predictor were entered into a single mixed model, directly testing whether the effect differed between timepoints rather than comparing two separately fit models. The type of forgetting among toddlers who formed associative bindings at initial retrieval was modelled with multinomial logistic regression (multinom; reference category: retained binding), covarying for retrieval delay, age, sex, baseline binding strength, and encoding trials. All nested model comparisons used likelihood ratio tests (LRT; anova). Trial within an encoding block was modelled as a categorical factor, because accuracy rose steeply from the first to the second trial of a context and then levelled off. Stability of encoding and retrieval performance (session 1 vs. session 2) was assessed with Pearson correlations. Paired categorical outcomes (e.g., recall of one context vs. another within the same session) were compared using McNemar's test. Multiple comparisons across jointly tested outcomes were corrected using the Benjamini-Hochberg procedure. The three continuous HMM outcomes (probability of full binding, location-only responding, and complete forgetting) were modelled and reported separately rather than as a Benjamini-Hochberg-corrected family, as they are complementary descriptions of the same compositional classification (the three probabilities sum to 1 for each child) rather than independent hypotheses. Within each of these models, the nine terms (timepoint, the four predictors, and their interactions with timepoint) were corrected using the Benjamini-Hochberg procedure, and adjusted P values are labelled as such. Follow-up contrasts and simple slopes within a single model were corrected using the Holm procedure, and adjusted P values are labelled as such. For the maternal COVID-exposure cohort comparison, related outcomes (e.g., accuracy and random error) were tested jointly with MANOVA (Pillai's trace) as an omnibus test, followed by per-outcome linear mixed-effects models with Benjamini-Hochberg correction.

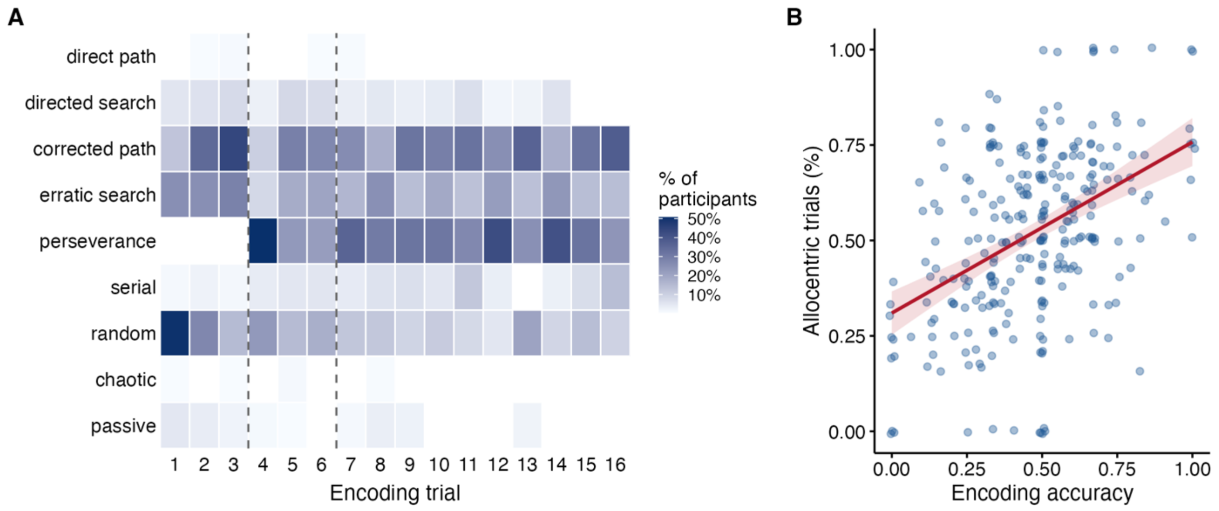


**Extended Data Fig. 1.**

**Search strategy detail and relationship to encoding accuracy.** (A) Proportion of toddlers classified into each of the nine individual search-strategy types (direct path, directed search, corrected path, erratic search, perseverance, serial, random, chaotic, passive) across encoding trials, shown as a heatmap; darker shading indicates a higher proportion of toddlers using that strategy on that trial. (B) Proportion of allocentric-strategy trials per toddler plotted against encoding accuracy in the same session; line, linear fit; shading, 95% confidence interval; points are individual toddlers. N = 270 toddlers (one point per toddler in B); toddlers per trial as in Fig. 2.

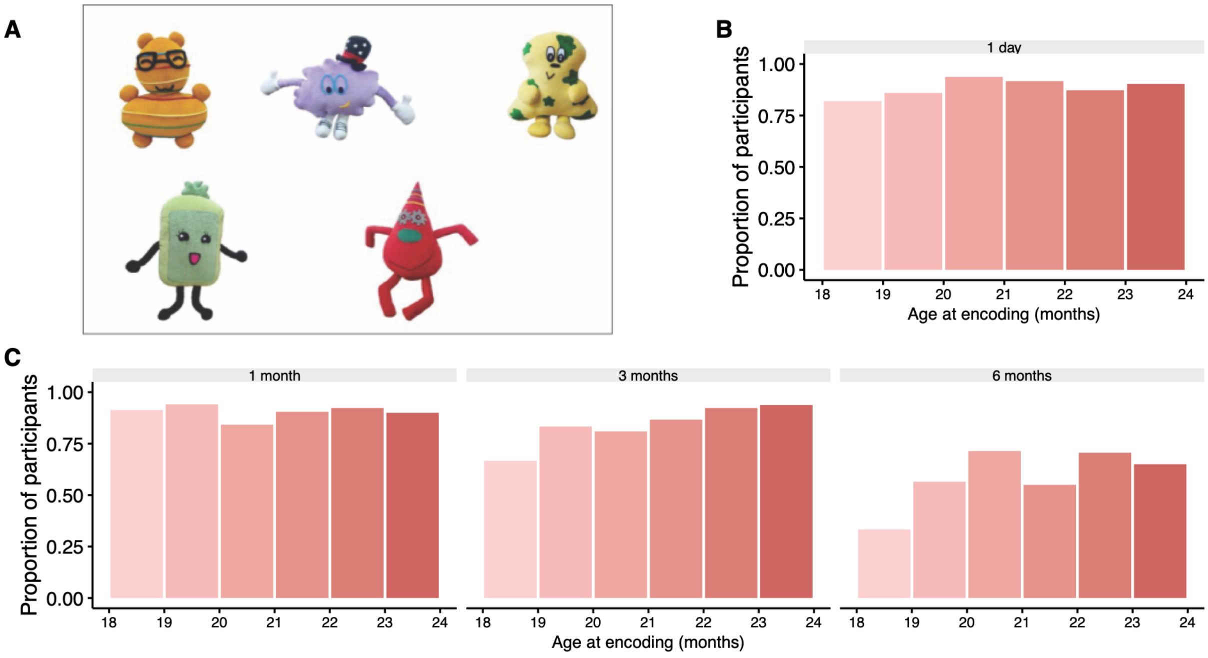


**Extended Data Fig. 2.**

**Item recall at initial and delayed retrieval.** (A) Schematic of the toy stimulus. (B) Proportion of toddlers who correctly recalled the toy at IR (1 day after encoding), by age at encoding (months); n = 355 toddlers. (C) Proportion of toddlers who correctly recalled the toy at DR, by age at encoding (months) and retrieval delay (1, 3 or 6 months; n = 113, 117 and 113 toddlers). Bars show proportions per one-month age bin, each toddler contributes one observation per panel.

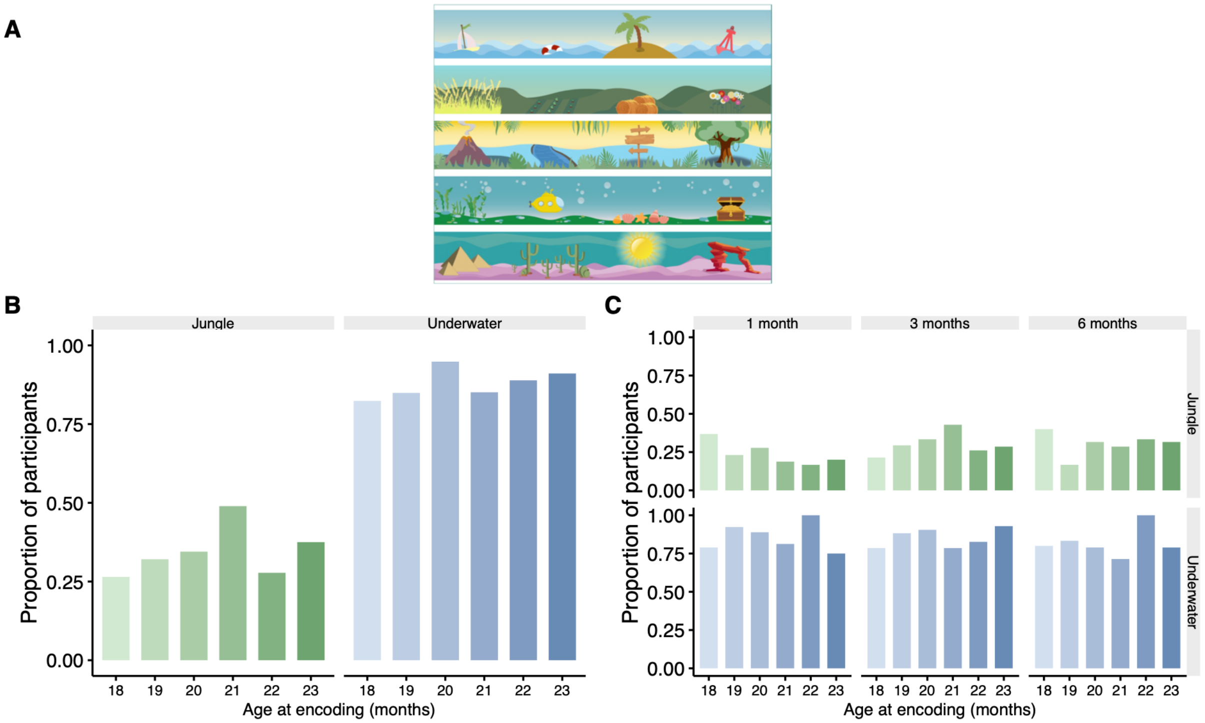


**Extended Data Fig. 3.**

**Context recall at initial and delayed retrieval.** (A) Schematic of the contexts used across encoding and retrieval sessions. (B) Proportion of toddlers who correctly recalled each scene at IR (1 day after encoding), by age at encoding (months); n = 302 toddlers, each scored on both scenes. (C) Proportion of toddlers who correctly recalled each scene at DR, by age at encoding (months) and retrieval delay (1, 3 or 6 months; n = 98, 103 and 108 toddlers). Bars show proportions per one-month age bin, each toddler contributes one observation per scene.

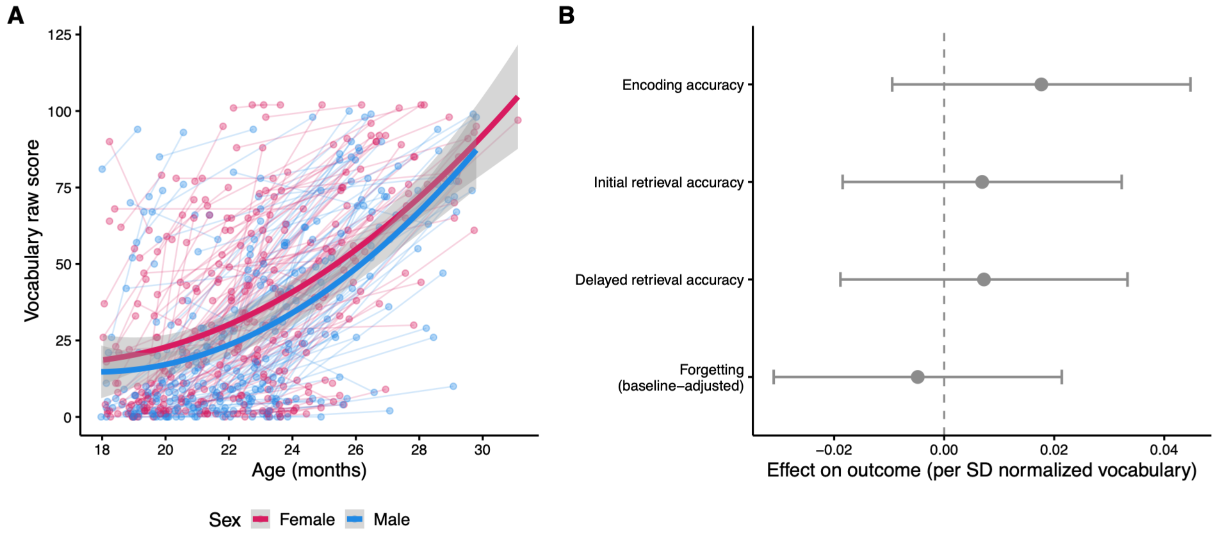


**Extended Data Fig. 4.**

**Language development and its relationship to memory performance.** (**A**) Vocabulary raw score (FRAKIS-K) by age at assessment (months), separately for males and females. Thin lines connect each toddler's time 1 (IR) and time 2 (DR) scores; bold lines show the quadratic fit by sex (linear age term insufficient, P = 0.003). n = 325 toddlers, 278 with both time points. (**B**) Vocabulary as a predictor of memory performance: standardised effect (points) with 95% confidence intervals (bars) of an age- and sex-normalised vocabulary score (raw vocabulary residualised on quadratic age and sex, then standardised) at encoding, on encoding accuracy, IR accuracy, DR accuracy and forgetting (IR minus DR, adjusted for IR accuracy). Linear models; P values Benjamini-Hochberg adjusted across the four outcomes; n = 321 toddlers.
